# Operations Near the Edge of cognitive decline: Behavioral and Neural Correlates of cognitive load in Mild cognitive impairment

**DOI:** 10.64898/2026.01.04.697587

**Authors:** Adrija Chatterjee, BS Shreelekha, Hari Prakash Tiwari, B Nandini Priyanka, Subasree Ramakrishnan, Avinash Singh, Pragathi Priyadharsini Balasubramani

## Abstract

**Motivation:** Early detection of cognitive strain is essential for identifying Mild Cognitive Impairment (MCI) and its progression to more severe neurodegenerative diseases. This study investigates cognitive strain in MCI by examining both behavioral and neural correlates during a real-world task.

**Methods:** Twenty-four older adults (M=69, 14 male, 10 female), including 11 with MCI and 13 healthy controls (MMSE >22/30), performed a spatial navigation task under varying memory load. We recorded brain activity using electroencephalography (EEG) and a mismatch negativity (MMN) paradigm to capture error-related potentials (ErrPs), which reflect cognitive load and processing errors.

**Results:** Both groups showed a decline in working memory performance as cognitive load increased (β = -41.39, SE = 10.73, t(90) = -3.86, p < .001). MCI participants, however, exhibited more efficient use of non-intrinsic (allocentric) reference frames under high cognitive load (β = 2.47, SE = 0.95, t(82) = 2.61, p = .009). MCI individuals also displayed larger ErrPs in central brain regions, indicating greater neural resource allocation. Additionally, MCI participants showed a decrease in parietal-to-frontal theta dominance under high load (β = -0.68, SE = 0.32, t(90) = -2.12, p = .037), and reduced fronto-parietal connectivity.

**Discussion:** These findings suggest that MCI participants experience greater cognitive strain, evidenced by increased neural resource allocation and strategy-shifting under load. This heightened strain may serve as an early marker of MCI progression, offering potential for early detection of pathological changes.

## INTRODUCTION

### Cognitive load and MCI

Mild Cognitive Impairment (MCI) represents an early stage of cognitive decline (Albert et al., 2011), often viewed as a transitional phase between normal aging and Alzheimer’s disease (AD) (Petersen et al., 2001). Individuals with MCI exhibit cognitive deficits that surpass what is typically expected from normal aging, although these changes may be subtle and not immediately impair daily functioning. In particular, MCI manifests through subtle memory deficits, visuospatial challenges, and other cognitive disruptions that are noticeable to the individual or their close social circle. Despite these challenges, individuals with MCI may still function relatively well in day-to-day activities, though the increased cognitive strain can be significant. Understanding the neural and behavioral markers of cognitive resources is crucial for identifying the onset of MCI, particularly for early diagnosis and intervention strategies aimed at preventing progression to AD.

A key aspect of cognitive function is the ability to manage cognitive load, or the mental effort required to perform tasks of varying complexity. For individuals with MCI, the ability to manage cognitive load becomes increasingly compromised. As task demands exceed the available cognitive resources, individuals with MCI often experience heightened cognitive strain. This strain, in turn, exacerbates memory decline and can accelerate the trajectory towards dementia (Schlosser et al., 2020). Working memory, in particular, is a central cognitive function impacted by this strain. Working memory relies on the ability to hold and manipulate information in the short term, but it also draws on finite neural resources. In the MCI population, this system can become overwhelmed, leading to difficulties in task performance and further cognitive deterioration.

Cognitive resource theory offers a framework for understanding these dynamics, suggesting that memory retrieval is not only influenced by the availability of relevant information but also by the competition for neural resources, such as neuromodulators, for the irrelevant information. Inhibitory mechanisms normally suppress such interference (Bjork & Bjork, 1996), but retrieval inhibition declines with aging. Older adults, including MCI, often struggle to suppress irrelevant stimuli and thoughts, reducing resources available for task-relevant processing and thereby accelerating cognitive decline (May & Hasher, 1998). This reduction in the ability to filter out irrelevant stimuli or thoughts diminishes the cognitive resources available for task-relevant processing. The result is an inefficient allocation of resources that further burdens memory systems, accelerating cognitive decline.

### Egocentric and Allocentric spatial navigation in MCI

Spatial navigation or spatial disorientation is one of the early symptoms in people with mild cognitive impairment (Cammisuli et al., 2024). There are at least two distinct computations in navigation: the egocentric frame of reference based processing, and the allocentric processing, which are relevant for characterizing Mild Cognitively Impaired (MCI) patients. The egocentric reference frame encodes spatial information relative to the self, whereas an allocentric reference frame encodes spatial relations between external objects, independent of the observer and reflects a preference for environment-centred, map-like spatial representation. The dominant viewpoint regarding MCIs can be generally divided into two main approaches: one group of research emphasizes the disorientation due to visuo-perceptual, and its connection with visual-spatial attention and optic flow perception. In contrast, another set emphasizes deficits in cognitive mapping in these patients, particularly regarding their use of allocentric navigation. Moreover, some studies suggest a possible link between spatial disorientation in individuals with Alzheimer’s Disease (AD) and MCI and the functioning of both the parietal and medial temporal lobes. Recent reviews in this subject either support the allocentric perspective, suggest integrating visuo-perceptual factors and cognitive mapping, or promote a multifocal theory of disease progression ranging from lateral to frontal and temporal to parietal brain regions, accompanied by related cognitive impairments.

Importantly, studies suggest the dominant role of the retrosplenial cortex (RSC) relative to frontal areas for allocentric spatial-cognitive map processing. Identifying the precise mechanisms mediating cognitive impairment can assist in engaging dysfunctional circuits and rehabilitating using smarter closed-loop stimulation paradigms (Cammisuli et al., 2024; Lin et al., 2015; Sulpizio et al., 2016; Zhang & Ekstrom, 2013).

Navigation also relies on the flexibility of switching between the reference frames of computation for effective navigation. Traditionally, it was believed that the brain’s spatial map is coded based on an allocentric frame involving the hippocampus and neighbouring regions (Wang et al., 2020). Detailed research shows primary upstream regions like the postrhinal cortex contain egocentric boundary cells (EBC) while downstream processing regions like the parasubiculum and medial entorhinal cortex contain conjunctive EBCs and head direction (HD) cells (Gofman et al., 2019). In particular, the HD cells assist in encoding the spatial context of an incident by converting egocentric sensory data into an allocentric map (Wang et al., 2020). The reverse process of transforming stored allocentric codes back into egocentric viewpoints enables the reconstruction of first-person perspectives during retrieval (Bicanski & Burgess, 2018). Since loss of episodic memory, due to dysfunctional hippocampal circuitry, is an early sign of Alzheimer’s disease (AD) (Weintraub et al., 2012), this frame switching that occurs during encoding and retrieval can be used as a cognitive indicator.

### Our study

Though many studies focus on virtual navigation-based understanding of neural processes, there are gaps in translating the scientific understanding to the real world (Kalantari et al., 2023; Li et al., 2021). In our study, we further probe the circuits using a passive cognitive task that hints at the resource distribution and perturbs the system by increasing the load on working memory, while participants navigate in real-world settings. This study investigates the relationship between working memory (WM) load and spatial reference frame use, using a real-world navigational task. We study two levels of experimental load through a working memory task (Hoppe et al., 2000), and two measures for the studying navigational orientation-egocentric and allocentric homing angle performance. Furthermore, we use electroencephalogram (EEG) to track the neurophysiological basis of cognitive load during navigation, and even administer a passive auditory mismatch negativity (MMN) paradigm (Näätänen et al., 2014) to measure global cognitive resource strain in terms of the magnitude of Error Related Potentials (ErrPs).

To quantify the cognitive strain and resource availability in MCI, auditory-based neurophysiological markers such as MMN provide valuable insight. MMN is an event-related potential (ERP) component elicited when an auditory stimulus deviates from a regular pattern typically occurring in a passive listening context without the need for active attention.

Reduced MMN responses may also indicate a decreased availability of cognitive resources to maintain and update sensory representations in working memory (Brima et al., 2019; John et al., 2024). Since MMN does not rely on overt task performance, it serves as a non-invasive, objective measure of underlying cognitive dysfunction—especially useful for populations where behavioral assessments may be inconclusive or limited by compensatory strategies.

We hypothesize that identifying the extent of ego-versus-allocentric performance changes, through real-time physiological signatures of mental strain during high cognitive load conditions, can offer suitable markers of clinical significance for cognitive training in MCI patients. By combining behavioral assessments of cognitive load in a navigational paradigm with neurophysiological measurements of evoked potentials, we can gain a more comprehensive understanding of the cognitive resources available to individuals with MCI. We propose that this approach not only provides insights into the cognitive challenges MCI patients face but also provides valuable tools for early diagnosis and the development of targeted interventions aimed at alleviating cognitive strain and potentially delaying the progression to Alzheimer’s disease.

*Our specific research questions include:* 1) How does general cognitive strain differ in MCI? For this, we assess the extent of resource allocation as indicated by ErrPs for unexpected auditory tones. 2) How does specific cognitive strain in navigation exhibit in MCI? For this, we assess the relative dominance of frontal and parietal activations and their connectivity, and investigate their relation to subjective spatial frame preferences.

## METHODS

### Participants

The study was conducted on a sample of 24 people (14 M and 10 F) above 55 years of age (M=69, SD=1.64). It was checked that none of the participants had hearing impairments, had been diagnosed with a neurological condition, or had ever exhibited behavioural problems. Mini-Mental State Examination, MMSE (Arevalo-Rodriguez et al., 2015) was used as a screening tool to segregate participants into healthy and mild cognitive impairment (MCI) categories. Participants with a MMSE score of more than 22 were considered healthy (Ganguli et al., 1995; Pezzotti et al., 2008). There were 11 MCI and 13 healthy participants, and the average MMSE scores for both groups combined (out of 30, M=22, SD=3.68). Participants with neurological disorders, hearing impairment, or conditions that could influence research investigation. Participants provided informed consent, received monetary compensation, and the protocol was approved by the Indian Institute of Technology Kanpur ethics committee.

#### Baseline assessments

Digit numbering task (DNT) threshold through psychometry. Participants were recited number sequences to recall after a 7-second delay, inspired by earlier studies such as (Stoet, 2010, 2017). The length of the sequence increases with each trial, starting from a one-digit sequence. Each sequence length was tested over three trials. In any subject, the lower threshold was the longest sequence recited with 100% accuracy (3/3 trials), and the upper threshold was the longest sequence recited with 0% accuracy (0/3 trials, Upper threshold-M=6, SD=1.71, lower threshold-M=4, SD=1.28).

Reference frame proclivity test (RFPT). This test was used to classify the participants as intrinsically egocentric or allocentric navigators (Gramann et al., 2005, 2010; Yang et al., 2021). Participants navigated through a tunnel on the flat-screen monitor that included direction changes of various angles in the horizontal plane. At the end, they indicated the tunnel’s turn: if their response matched the direction of the homing vector computed with respect to the participant’s heading direction. The intrinsic navigational preference for each participant was determined by the proportion of egocentric versus allocentric responses out of 20 trials: a proportion greater than 0.5 (egocentric responses) indicated an egocentric preference, while less than 0.5 indicated an allocentric preference.

EEG Baseline. Participants were seated and instructed to remain idle with eyes open for five minutes, followed by five minutes with eyes closed, while an auditory oddball mismatch negativity (MMN) task was administered throughout the study.

Pre-Trial Instructions. Most participants were unfamiliar with the devices used in this experiment; therefore, a detailed demonstration was provided on how to use the Anglo-Meter app before the actual task (see Figure 1). The Anglometer consisted of two 180-degree protractors joined together, forming a full 360-degree scale. Participants were shown how to mark egocentric and allocentric angles on the app. It was explained that they were positioned at the origin, and with the area above the origin corresponding to forward directions and below indicating behind them. To help them grasp the task, we used objects in the room for hands-on practice. For example, we placed objects in different quadrants and showed participants how to translate the object’s position into a protractor measurement on the app. For instance, if an object (Object 1) was directly behind them (180°), they were asked to identify the position of a second object (e.g., a table) relative to Object 1, treating Object 1 as the new reference point (0°). This practical demonstration ensured participants were comfortable using the app and understanding the task.

**Figure 1:**
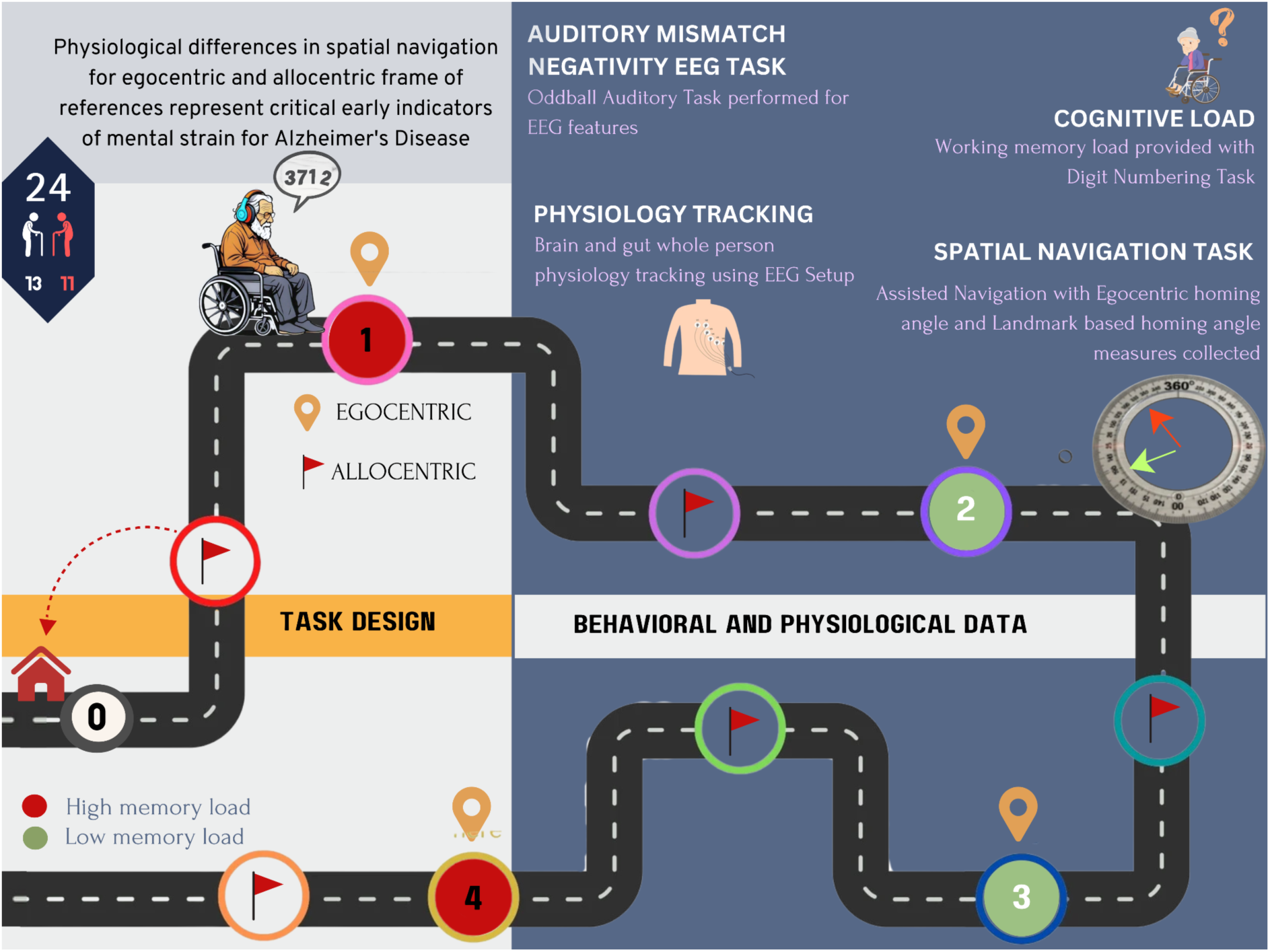
Study illustration. The figure illustrates the spatial navigation paradigm design probing egocentric and allocentric navigation abilities under two levels of cognitive load with simultaneous physiology tracking (EEG) and passive Mismatch negativity task in the background. The illustration also shows landmark locations in each trial that could be low or high working memory load, and the measurement of errors using the anglometer.

Practice trial. Further, a practice (control) trial was administered before the experiment. The memory load (digit length) was set to 3, and one turn involved a navigation route of approximately 50 meters, which was kept consistent for all participants and was chosen for navigational load. Ego-centric and allo-centric response acquisition, landmark recognition, was also performed during the control trial.

#### Main Experiment Design

Participants navigated a real-world environment while seated in a wheelchair to accommodate elderly individuals with mobility limitations. The navigation speed was deliberately set to match a typical walking pace (Average: 1.51 km/hr, 0.77-2.10km/hr). Participants were asked to memorise the digit sequence in the presented order (lengths 3-8) depending on their baseline thresholds, with digits delivered one at a time, approximately equi-spaced for the length of the trial. They were informed beforehand about two landmarks that would appear along the path during the start of the trial, and instructed to attend to them while navigating. The paradigm followed a 2×2 factorial design: Working Memory Load: High vs. Low Load; Navigational Load: High vs. Low Load, as explained below.

Working memory load manipulation. The Digit Numbering task (Baddeley, 2012; Baggini & Ricciardelli, 2025) was used to induce load on the working memory capacity of the participant. *High-load trials:* Sequences with one digit less than their Digit Numbering task, DNT baseline upper threshold. *Low-load trials:* sequences with digits equal to or fewer than the DNT baseline lower threshold are classified as low-load trials. There were four main trials: two high-load trials, two low-load trials. The order of the trials was kept fixed - high, low, low, high (Figure 1).

Navigational load manipulation. The home location was constant across all trials. Each trial required three turns for completion. Consequently, the number of turns increased cumulatively across trials as the trials occurred in a continuous manner from the start position, adding to the cognitive demand by requiring continuous path recalibration. The first two trials were classified as low-navigational load trials, and the next two are high-navigational load trials **(Figure 1)**.

The average length of each trial was 67.5±22.4 meters covered in approximately 3.75±1.26 minutes with about 3 turns *that kept accumulating with every trial*; the participants had wireless headphones on, and EEG was recorded simultaneously.

Mismatch Negativity Paradigm. A passive auditory version was administered via wireless headphones, consisting of a sequence of standard tones (1000 Hz, 100 msec duration) presented at an inter-stimulus interval of 900 msec. Rare deviant tones (1000 Hz, 180 msec duration) occurred 20% of the time, randomly distributed between 10% and 90% of the trial length, with a stimulus onset asynchrony of 450 msec.

##### Outcome measures

Primary outcome measures were collected post-trial **(Table 1)**. Further, on a scale of 1-4, difficulty ratings on remembering the sequence and making spatial decisions were also collected.

**Table 1:**
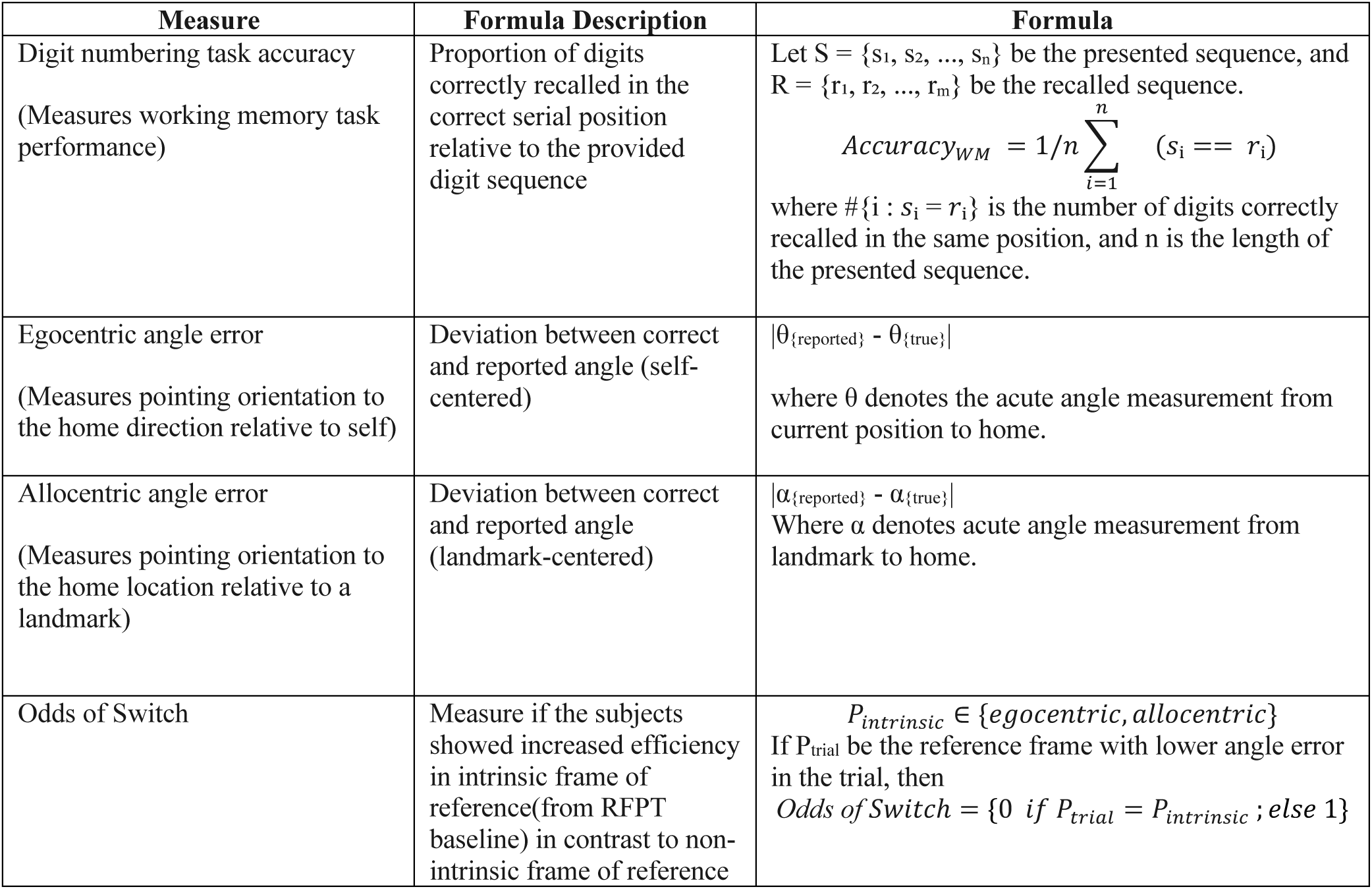
Post-trial outcome measures and their formulation used in the study. The table presents the formula and its description of the DNT task, Spatial navigation errors in different frames of reference, odds of switch in any trial.

#### Physiology methods

##### Electroencephalography (EEG)

Electroencephalogram (EEG) was recorded using standard 10-20 system (Klem et al., 1999; Nuwer et al., 1998) montage with 11 recording electrodes (Fp1, Fp2, F3, F4, C3, C4, P3, P4, Cz, Fz and Pz), 1 reference, 1 ground electrode, manufactured by Clarity medicals in the name of BrainTech machine. The data was stored as text files with timestamps from standard clock time in 256Hz resolution. For about 3 out of 24 people (M2, M3, F4), EEG data was acquired using another device- a 24-channel SMARTING cap with a semi-dry and wireless electrode layout (Next EEG—new human interface, MBT) and in these specific cases, data were acquired at 500 Hz sampling frequency in 1 participant (M3), and 250 Hz in the other two. In all participants, event markers were integrated using LSL and data files were stored in xdf format. Data was imported to MATLAB (The MathWorks Inc., 2022) and processed within EEGLAB v2023.1 (Delorme & Makeig, 2004).

Data Preprocessing. For each file, the data were imported and downsampled to 250 Hz to ensure alignment with stimulus timing. Approximately 5 seconds of flat signal were trimmed from the end of each recording to remove artifacts. Subsequently, the data were filtered using a zero-phase FIR filter with a passband of 0.5–40 Hz and a filter order of 9000.

Artifact-prone channels were detected and removed by executing the *clean_channels function*. Independent component analysis (ICA) was run (*pop_runica)*, and the components were automatically labelled using *ICLabel*. Components less than 5% probable as neural and >80% probable of eye, heart, line noise and muscle were rejected and eliminated using *pop_subcomp function.* Further non-stationary artifacts were cleaned by executing *clean_rawdata function.* Channels with flatline periods longer than 5 s were removed, and channel reliability was assessed by correlation within the 0.25–0.75 Hz band, with those falling below a correlation threshold of 0.8 or exceeding 4 standard deviations of noise rejected. Artifact Subspace Reconstruction (ASR) was applied with a burst detection threshold of 5 SD, using the entire recording as the analysis window. Eliminated channels were interpolated using *pop_interp function* employing spherical splines, and the data were average-referenced (using *pop_reref function)*. Preprocessed EEG datasets were stored for further analysis.

##### ERP Segmentation and Mismatch Negativity (MMN) Computation, ERPdiff

For computing mismatch negativity, EEG trials were epoched relative to auditory stimuli, which was present for about 2.64 mins±19.5 secs of any trial, designated as standard and deviant with a window from -500 msec to +1000 msec relative to stimulus onset. Baseline correction was applied, and baseline was computed as an average of the pre-stimulus interval (-500 msec to 0 msec). High-amplitude artifact epochs were rejected using EEGLAB’s (using *pop_autorej function),* and a maximum rejection of 10% in any subject to compute the event related potentials (ERPs). The MMN waveform (ERPdiff) was calculated as the deviant ERP minus the standard ERP along with the following steps, for each channel. ERPs were truncated to a post-stimulus interval equal to 100–400 msec to separate the MMN component. Channel-wise MMN waveforms were normalized by mean subtraction across time.

MMN Peak Detection and Regional Analysis. To determine the MMN peak, the difference waveform in each channel was searched for the five most negative local minima. For each minimum, the mean amplitude within a **±**10 sample point window was computed, and the minimum of these local averages was saved as the peak MMN amplitude. Electrodes were classified into frontal (Fz, F3, F4), central (Cz, C3, C4), and parietal (Pz, P3, P4) regions. Regional MMN values were calculated as the mean of constituent channel amplitudes.

##### Generalised eigenvalue decomposition (GED), EEGTopo

We employed Generalized Eigen Decomposition (Cohen, 2022) to identify spatial sources that orthogonally differentiate neural activity in the experimental condition from the reference (eyes-open resting state part of the baseline trial). To account for how activation pattern changes with respect to experimental conditions (different working memory load), the EEG recordings were segmented into trials, and GED was applied on a trial-wise basis, each time using the eyes-open baseline as the reference. Component map (Compmap) was obtained from each trial as Compmap= w1TCs, where w1T-the first eigenvector(with the highest eigenvalue) that captures the most task-specific variance relative to baseline. Cs-signal covariance matrix represents the spatial covariance structure of the EEG data during any trial. It captures how EEG signals from different channels co-vary across time. This matrix is used in GED to find spatial filters (eigenvectors) that maximize task-specific variance (CovS) relative to baseline variance (CovR).

EEGTopo. Two primary areas of focus were the frontal and parietal cortices. First component maps with max eigenvalue were extracted from specific electrode sites—Fz, F3, and F4 for the frontal region, and Pz, P3, and P4 for the parietal region. For each trial, the maximum absolute value across these sets of electrodes was calculated to represent the dominant activation within each region. The difference between frontal and parietal values was then computed. Based on the distribution of these score differences (in percentiles), each trial was categorized into one of three region labels: Frontal-dominant, Parietal-dominant or otherwise.

##### Statistics

We applied a generalized linear regression model to predict group membership (MCI = 1, Healthy = 0) using performance measures including working memory accuracy, egocentric and allocentric angle errors, and switch performance, along with their interactions. All predictors were median split to transform to categorical variables. The model was implemented in MATLAB using the *fitglm* function. An 80:20 train–test split was performed using *cvpartition* function with a fixed random seed, and the model was trained on the training set and evaluated on the held-out test set. Based on the test set, Model AUC, sensitivity and specificity was computed. Finally, a K-fold (k=3) cross-validation was done to further assess model generalizability. Further, Bonferroni corrections were applied for multiple comparisons appropriately.

## RESULTS

In the study, healthy (N=13, MMSE 24.46±1.45) and MCI (N=11, 19.09±3.39) participants and MCI participants used assisted navigation conducted alongside continuous presentation of digit sequences and increasing navigational complexity, which together contributed to cognitive load. Through the task, participants executed an auditory oddball MMN task in a passive fashion, and concurrent EEG recordings were captured.

### 2.1 Behavioural feature results

Firstly, we asked how memory-based behavioural performance differed between participant groups. There was a significant main effect of working memory load on accuracy, with participants performing better in low-load trials compared to high-load trials (working memory load condition) *β* = -41.39, *SE* = 10.73, *t*(90) = -3.86, *p* < .001 (see **Figure 2a**). There was no significant effect found across navigational load trials. No significant interaction effects were seen between the group and the experimental load conditions.

**Figure 2:**
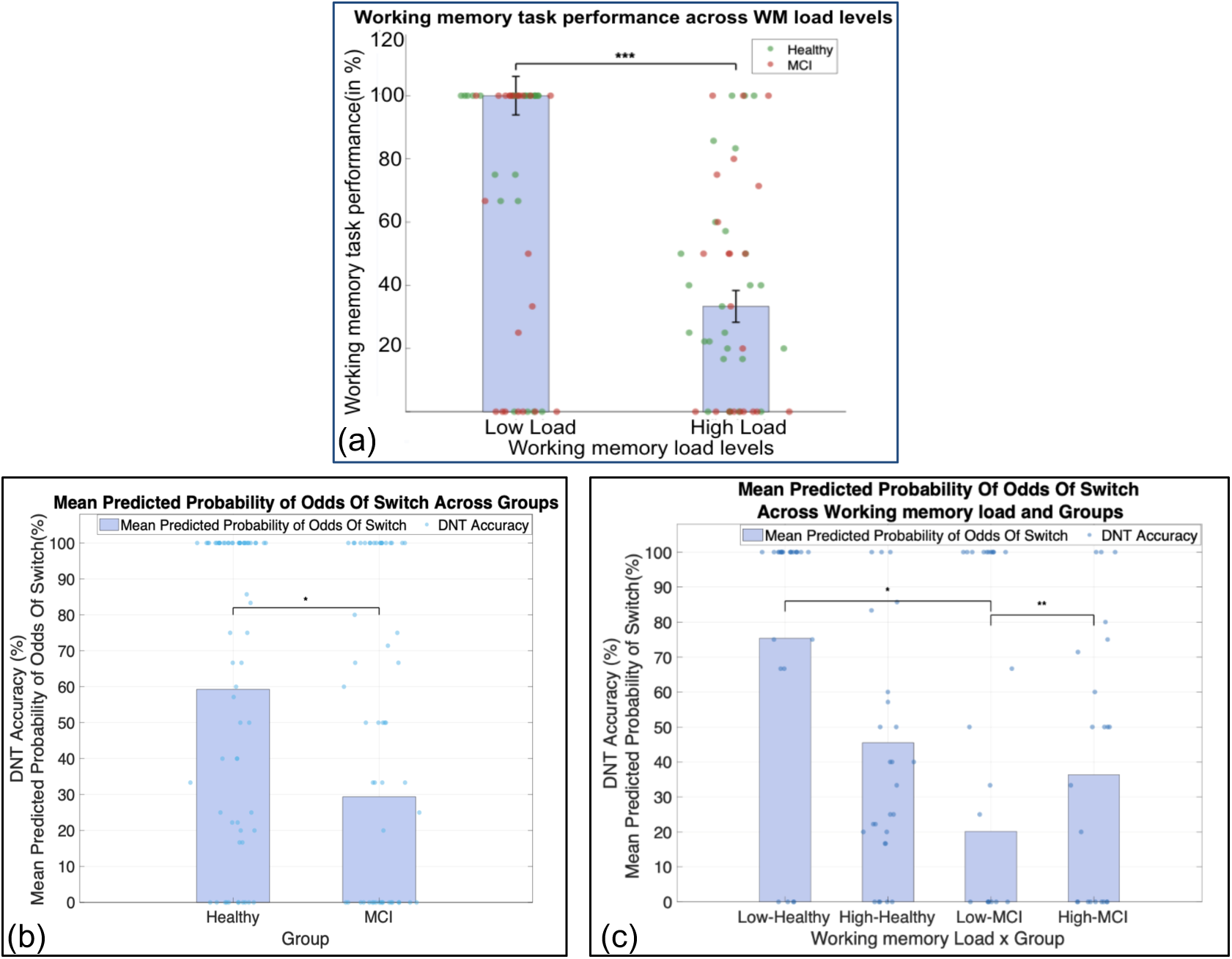
Behavioural feature results. (a) Performance in the Digit Numbering task (working memory profile task) across low and high working memory load. The scatter points show the trial results in participants. The bars represent the mean and the standard error. (b) Mean predicted odds of switch probability across groups (c) Across group x working memory load conditions.

Secondly, we asked whether their preferred frame of navigation (egocentric versus allocentric) changed with different memory loads. For this, we initially computed their intrinsic preferred frame of reference using the RFPT described in the methods. We later checked their current frame of reference (as described more in the methods). The switch to non-preferred frames of reference for spatial orientation, i.e. *odds of switch* was less in MCI in general (N=88, df=82, estimate= -1.869, SE= 0.83, tStat= -2.25, pValue=0.024). However, a significant interaction was also found between the group and working memory load condition. *For high working memory load trials, odds of switching were more frequent in MCI, while for low working memory load trials, the healthy cohort had more odds of switch than the MCI* (Group_MCI: working memory load interaction: N=88, df=82, estimate= 2.467, SE= 0.95, tStat= 2.61, pValue=0.009, **Figure 2b,c**).

#### 2.1.1 Prediction Model And Validation

We trained a predictive model on all subject trial-level data and analysed the importance of various features explaining the group status (MCI versus controls). A generalized linear regression model with behavioural predictors from memory tasks such as the recall with order accuracy, ego- and allo-centricity error, odds of switch, and their pairwise interactions, was developed to predict the probability of MCI. The model had a Fstat(63) = 2.59, p=0.004, AUC=0.87.

When the model was trained on 80% of the data and evaluated on the remaining 20%. The test set performance yielded an AUC of 0.722, a sensitivity of 71.43 %, a specificity of 77.78% and an accuracy was 66.67%. The optimal classification threshold was selected using Youden’s J statistic, which was 0.680.

To further assess the model’s generalization*, we used k-fold cross-validation.* The dataset was divided into 3 equal folds, with the model trained on K-1 folds and tested on the remaining fold. The average metrics across the folds were: AUC 0.738±0.04, sensitivity 70.85%, specificity 79.53%, and accuracy 67.50%. (see Supplementary material for detailed statistics on the features used in the model).

### 2.2 Neural Feature Results

We compute several *physiological features* to assess the eventual navigational and memory performance in our subjects. In EEG, we focused on mismatch negativity representation at the level of event-related potentials (ERPdiff), frontal or parietal dominance as assessed by topographically patterns obtained through Generalized EigenValue Decomposition (EEGtopo).

#### 2.2.1 ERPdiff: The MCI had differentiable mismatch negativity potential in the central, frontal and parietal regions

The peak difference between deviant and standard baseline corrected ERP signal (referred here as ERPdiff) within time units of 100-400 msecs is significantly different in the MCI: We specifically looked at three separate regions, Frontal, Central and Parietal, and found them all to be differentiating the MCI from Healthy with greater standard versus deviant activity in MCI population (frontal, *t(26) = –2.06*, *p < .05*, central, *t(26) = –4.54, p < .001*, and parietal regions, *t(26) = –2.41, p < .05*, **Figure 3**).

**Figure 3.**
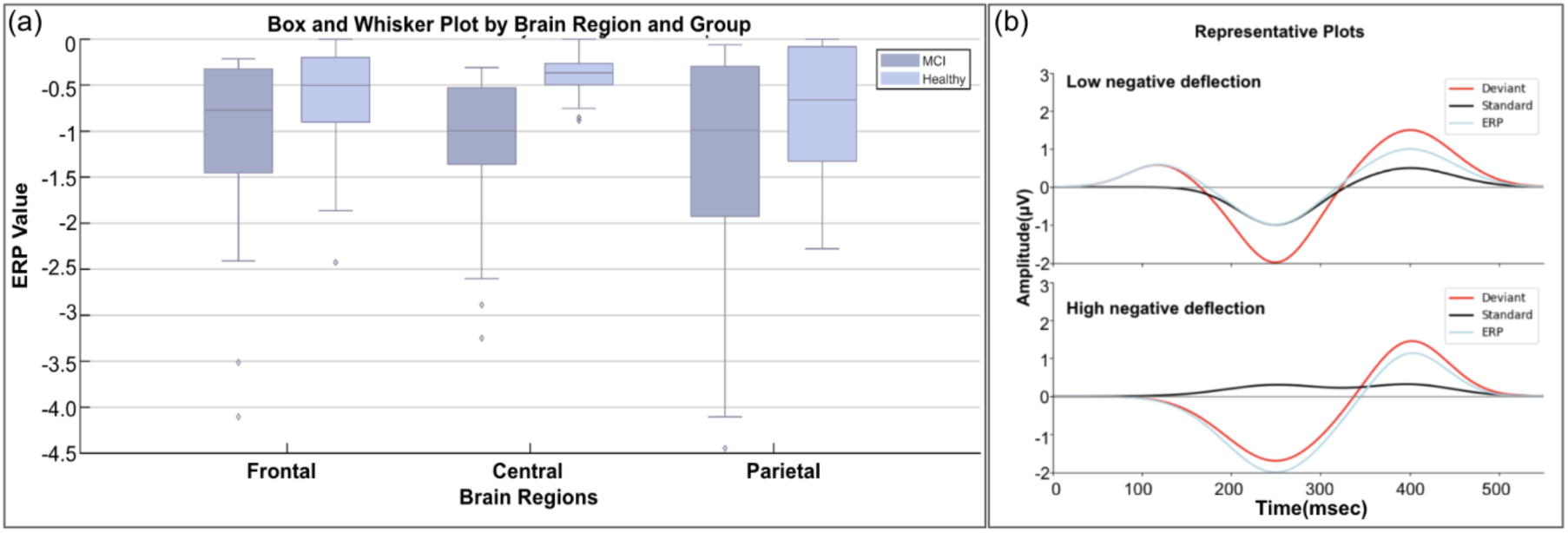
a) Difference between Deviant and Standard ERP signal (ERPdiff-within time units of 100-400 msecs), across Frontal, Central and Parietal regions in Healthy and MCI cohorts. The inset shows ERP representation plots for standard and deviant tone presentations at 0 msecs, and their differences (deviant-standard).

**Figure 4.**
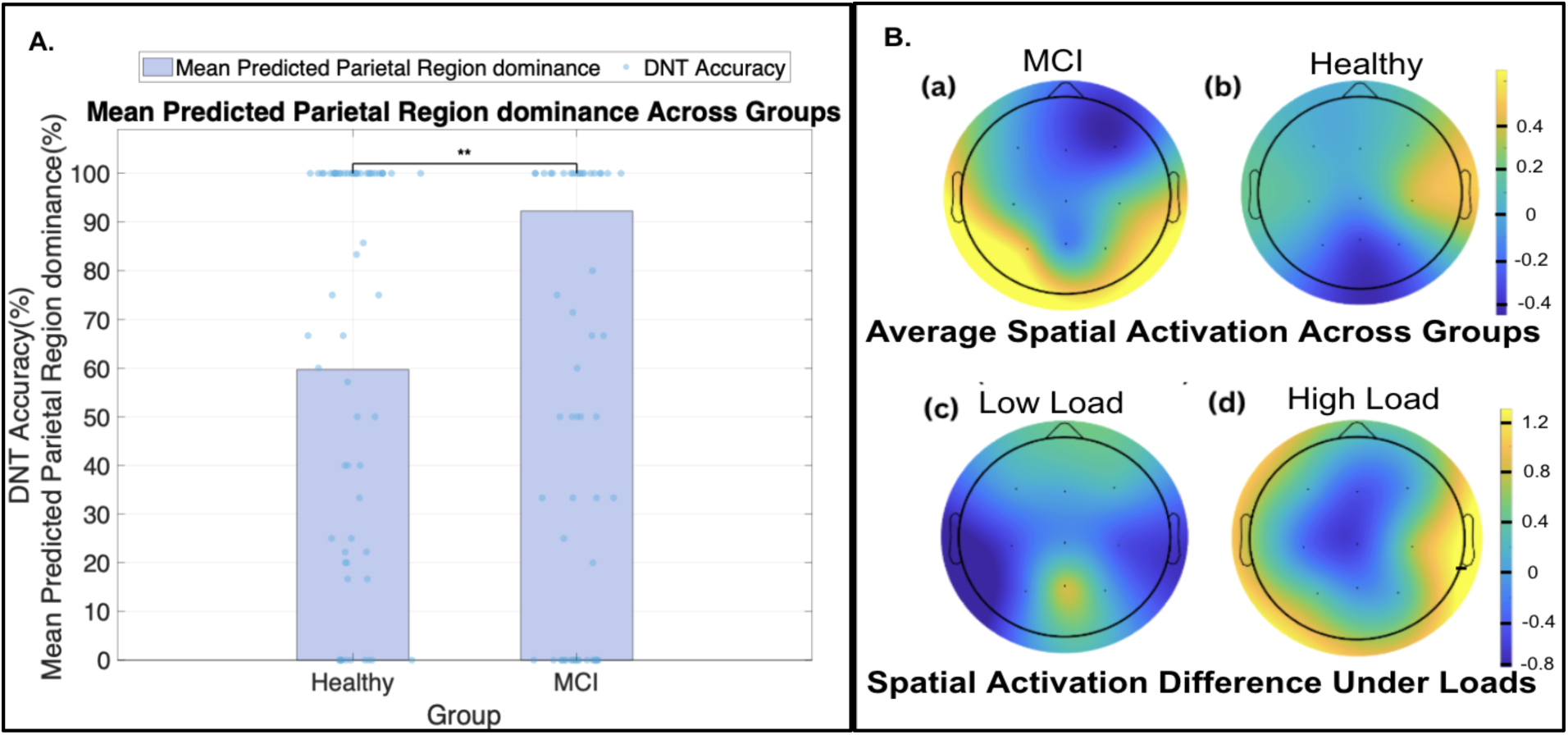
**A**. Mean predicted parietal region dominance (relative to frontal) in theta, derived from GED across healthy and MCI. **B**.(a)Average topoplot for (a) MCI (b) Healthy (c) Spatial activation difference between MCI and Healthy under low load (d) high load.

Controlling for multiple comparisons, linear regression models with ERPdiff as the dependent variable for each region (Central, Parietal, and Frontal) and experimental conditions such as working memory load and navigational load as independent variables were performed. It was found that ERPdiff in the Central region was sensitive to the experimental load conditions, *with R² = 0.32 and p* = 0.019. We also found the *switch* to be *positively* correlated with EEGdiff in the parietal region, *β =* 0.63, *SE =* 0.29, *t(44) =* 2.17*, p = .*033.

#### 2.2.2 EEGtopo: Overall parietal to frontal dominance was reduced in the high load condition in MCI

Parietal dominance relative to the frontal activations was seen in MCI (*β* = 0.81, *SE* = 0.28, *t*(90) = 2.92, *p* = .004. However, this dominance was reduced during high working memory load conditions (significant negative interaction between working memory load and group: *β* = –0.68, *SE* = 0.32, *t*(90) = –2.12, *p* = .037) and the observed difference frontal-parietal activations were positively correlated to egocentric angle error (*β* = 0.55, *SE* = 0.19, *t*(70) = 2.91, *p* = 0.004).

#### 2.2.3. A decreased fronto-parietal connectivity correlated with increased errors

We performed connectivity analysis to gain a better understanding of the parietal to frontal shift with an increase in navigational error. The error variable was correlated with decreased frontal-theta to parietal-beta coupling, *β* = -7.12, *t(46)* =2.06, *p* = 0.001(as well as fronto-parietal theta coupling, *β* = -2.91, *t(46)* =1.17, *p* = 0.016).

## DISCUSSION

Alzheimer’s- the most common form of dementia (∼75%) is a progressive neurodegenerative disease (World Health Organization: WHO & World Health Organization: WHO, 2025);. Age is one of the primary risk factors for AD. Five per cent of people aged 65 to 74, 13.2% of people aged 75 to 84, and 33.4% of people aged 85 or older have Alzheimer’s dementia (2024 Alzheimer’s Disease Facts and Figures, 2024). Human life expectancy has increased over time, leading to an absolute increase in the number of people with AD. This number will rise without developing early prediction tools or efficient treatments (Winblad et al., 2016). Between 2000 and 2021, reported deaths from AD increased by more than 140% (2024 Alzheimer’s Disease Facts and Figures, 2024). According to the *Global Burden of Disease Study (GBDS) 2019,* from 2019 to 2050, the number of dementia cases will undergo a whopping increase of 166%, impacting the lives of ∼152·8 million individuals; these estimates are similar to those predicted by WHO (Ray et al., 2023). The greatest increase will be seen in countries with low socio-demographic index (Nichols et al., 2022), like India. In 2015, the global cost of dementia was estimated at US$818 billion, a figure that is expected to rise as the prevalence of dementia increases. Approximately 85% of these costs are associated with family and social care rather than medical care (Livingston et al., 2017).

Signs of wandering often surface in Alzheimer’s disease. However, they may be unable to retrace their steps and may become lost even in familiar surroundings; compromised object-based spatial navigation and spatio-cognitive map estimation-based navigation represent critical early indicators of mental strain for Alzheimer’s Disease (Cammisuli et al., 2024). Earlier studies also suggest that increased stress or mental load can aggravate the cognitive and memory decline in people (Schlosser et al., 2020). The disease progression is known as the AD continuum because there is no clear distinction between one stage and the other. It starts with unnoticeable brain changes *(Preclinical stage) and progresses* to cognitive functions that are impaired but not enough to interfere with daily life *(Mild Cognitive Impairment)* and ultimately *significant dementia* (Sperling et al., 2011; Albert et al., 2013).

The MCI stage is a transitional state between normal ageing and dementia, characterised by cognitive decline greater than expected for age but not meeting the criteria for dementia (Petersen et al., 2001). They show cognitive deterioration and maintain largely intact daily activities (Winblad et al., 2004). The older population can be screened for subtle cognitive impairments, such as difficulty with memory, language, and thinking. Screening tools like the *Mini-Mental State Examination (MMSE) and the Montreal Cognitive Assessment (MoCA)* are commonly used to test for MCI. Mild Cognitive Impairment (MCI) is a significant predictor of Alzheimer’s Disease (AD), with conversion rates varying across studies. Geslani et al. (2005) reported conversion rates of 41% after one year and 64% after two years in a clinical setting and Fischer et al. (2007) found conversion rates of 48.7% for amnestic MCI and 26.8% for non-amnestic MCI after 30 months in a community-based cohort.

### Our study

In this study, we measured cognitive decline across multiple cognitive functions, including auditory processing, memory, spatial navigation, and cognitive flexibility. Importantly, just their navigational performance under cognitive load in different frames of references (egocentric, allocentric error) were able to explain MCI status significantly. Measures like ErrP reflect the general capacity of the brain to manage cognitive load, irrespective of the specific task. We observe that MCI may be suffering from global neural inefficiency, highlighting when an individual is struggling with basic tasks, it can be an early indicator of general cognitive decline in MCI. On the other hand, the frontal dominance observed in the navigation task indicates a targeted compensatory strategy specific to spatial memory and navigation under higher cognitive load.

Using both ErrP and parietal-frontal tradeoff markers enables a multi-faceted understanding of MCI progression. Our study suggests that if a MCI subject shows increased cognitive load (ErrPs) but also demonstrates a spatial compensatory strategy (frontal dominance), it suggests that they might still have neural reserve or adaptive coping capacity, which could influence their long-term prognosis. These results further support that MCI participants further support the notion that they are operating near their cognitive limits even under task demands that exert lesser resources in healthy controls.

### Retrosplenial cortices involvement in MCI

In MCI, egocentric navigation, which relies on body-centred cues, may be impaired, prompting individuals to depend more on allocentric strategies supported by the hippocampus, often less affected in non-amnestic MCI. This is observed by the differential performance of MCI in egocentric and allocentric tasks for the same trials.

As they become more aware of their limitations, such as memory lapses, MCI individuals might increasingly use external cues in high-conflict scenarios to enhance discernment of relevant information, potentially improving navigation performance by focusing on distinct landmarks. Higher external sensory processing is observed with the higher ERP represented in MCI population compared to healthy around the central region. This compensatory mechanism, indicating reliance on preserved allocentric navigation, serves as a marker of amnestic MCI.

Early studies suggest the navigational circuits in the parietal region, including the Retrosplenial cortices and their interactions with the hippocampal circuit, contribute to an allocentric frame of reference-based navigation (Clark et al., 2018). The role of the hippocampus has been documented in many allocentric navigation studies that involve the importance of the hippocampus in spatial deficits in MCI and AD. In setups testing using Morris water mazes, the AD group with more defined symptoms faced challenges navigating using both hometown and wall cues, whereas the amnestic single-domain MCI group showed impairment only in tasks involving allocentric cues (Hort et al., 2007), indicating a peculiar deficiency indicates the hippocampus. Patients with AD also had difficulty navigating in a maze created in a virtual reality arena using a map, which required the translation of the allocentric view on the map to the maze in egocentric direction (Morganti et al., 2013).

Egocentric memory enables the recollection of one’s spatial position in relation to one’s own location and orientation in space. Structures in the inferior and superior parietal lobes showed increased activity during various egocentric tasks, as seen in the Morris water maze, Virtual City; (Parslow et al., 2004; Wolbers et al., 2004), as well as in an environment without characteristic objects (Wolbers et al., 2008).

### Cognitive flexibility is compromised in MCI

Low load conditions allow individuals to perform tasks more efficiently, with minimal cognitive strain. In control subjects, we observed the ability to switch between frames of reference, more as a sign of cognitive flexibility: When task demands increase, non-intrinsic frames may become more taxing due to the additional cognitive effort needed to process them. Therefore, in contrast to the MCI group, which shows more strain and compensatory behaviors driven by frontal and parietal interactions (Babiloni et al., 2006; Guarino et al., 2020), controls showed the ability to switch between reference frames without experiencing significant cognitive bottlenecks, allowing them to demonstrate better efficiency in non-intrinsic frames even at low load. Notably, we also observed fronto-parietal coupling with increased load in MCI.

### Real-time passive physiological markers can identify MCI

The poorer allocentric performance in MCI suggests RSC-driven parietal activity to be affected in our sample, offering a potential biomarker for closed-loop stimulation. Furthermore, the study asks whether an increased energy factor observed in the parietal area represents either compensatory mechanisms in MCI or inefficient resource management. The frontal dominance of power with increased cognitive load is an added sign of struggle observed in the MCI participants of our study. Our future work involves further validating our observations using high-resolution source localization.

These assessments are cost-effective and easy to administer. They provide a behavioural foundation for understanding the disease. Correlating behavioural findings with neurophysiological or electrophysiological measures enhances understanding of why some individuals demonstrate resilience while others do not. This approach supports the development of predictive models for Alzheimer’s disease. In particular, we propose applications for early detection using digital spatial navigation tasks that can track subtle changes in frame-of-reference usage and cognitive load handling, identifying early cognitive decline in MCI before overt symptoms appear. We predict that tools designed to promote flexible spatial strategies could help alleviate cognitive load and delay progression in MCI. One can also develop passive EEG systems or their related biosensors to monitor error potentials in real time to adjust task difficulty or provide feedback on cognitive strain.

Altogether, this study provides insight into the compensatory mechanisms that MCI individuals may employ under cognitive load, with parietal versus frontal tradeoff playing a prominent role. The hypothesis of RSC-driven parietal involvement suggests that early-stage cognitive decline may be marked by shifts in spatial processing strategies. These findings support the development of cognitive tools for early detection and personalized intervention in MCI.

### Limitations

Some of the limitations of our study include its small sample size and use of a low-density EEG system. Future work will be directed towards validating our current findings using high-resolution source localization methods, and testing them in clinical settings.

## Acknowledgements

to all the participants.

## Data availability statement

The data is available on reasonable requests to corresponding author.

## Supplementary Material

**Predictors of the behavioural GLM in the k-fold cross-validation**

**Figure.**
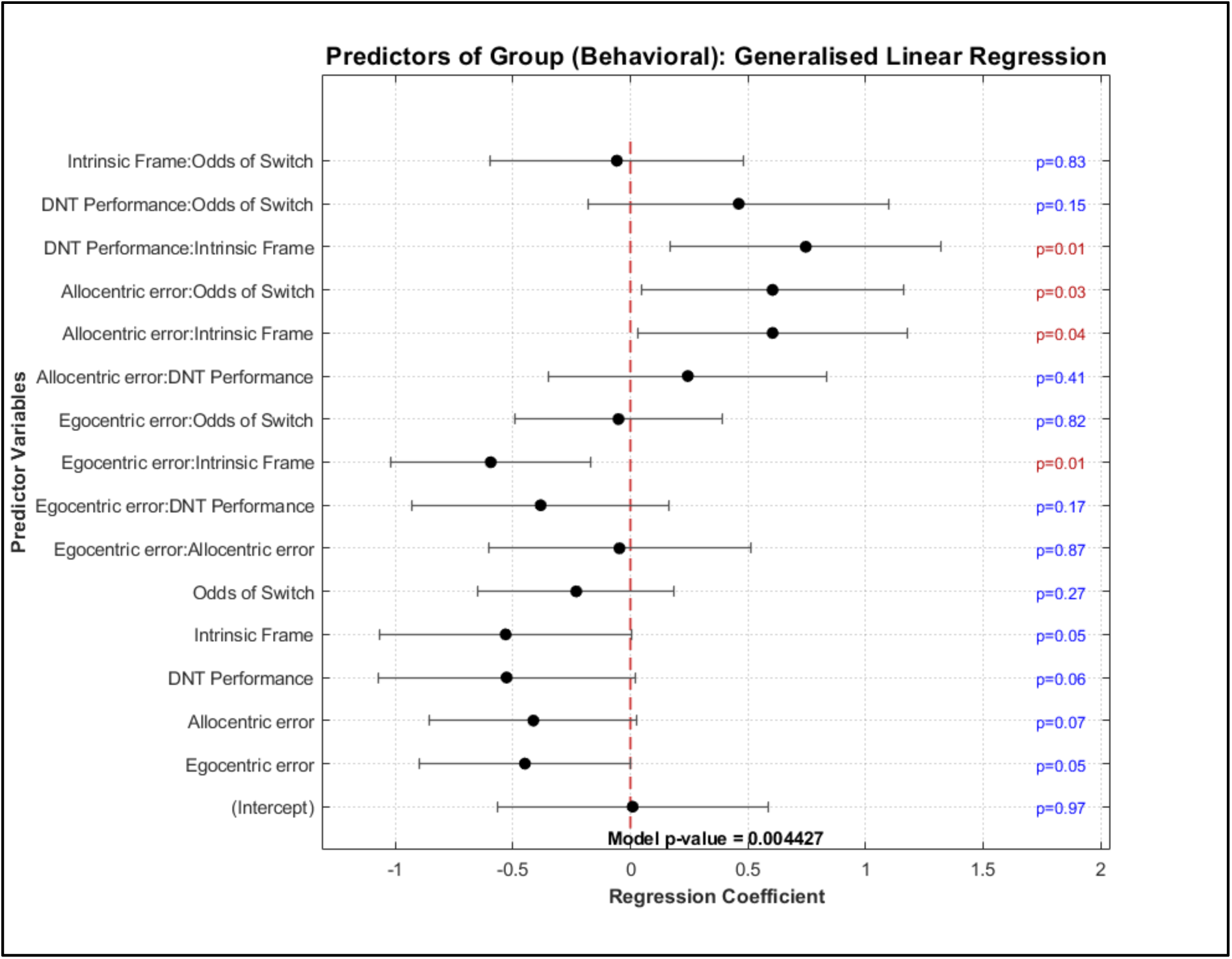
Base model and its estimates to predict the MCI status.

**Figure.**
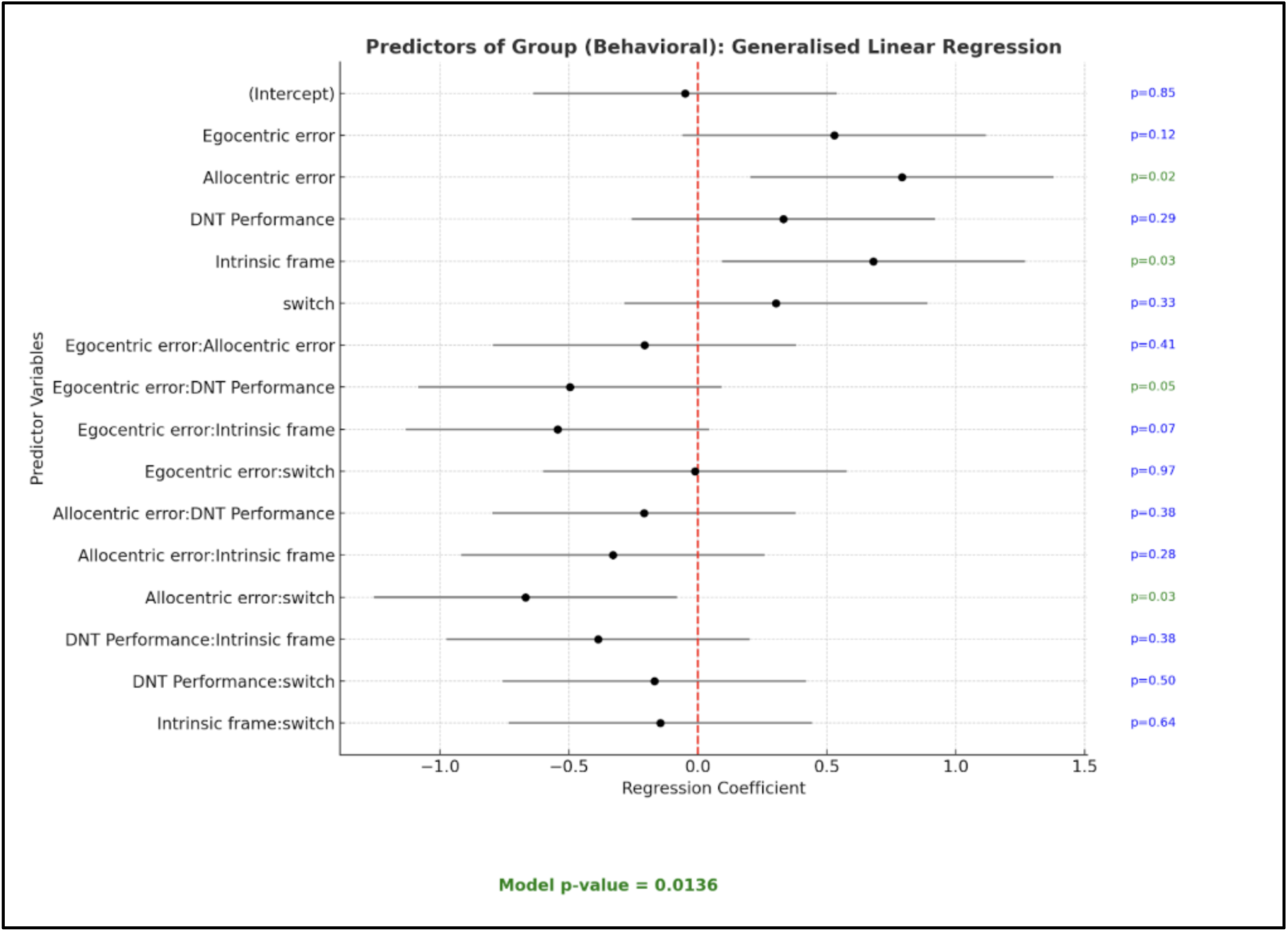
The model stats from 80:20 train-test split data.

**Figure.**
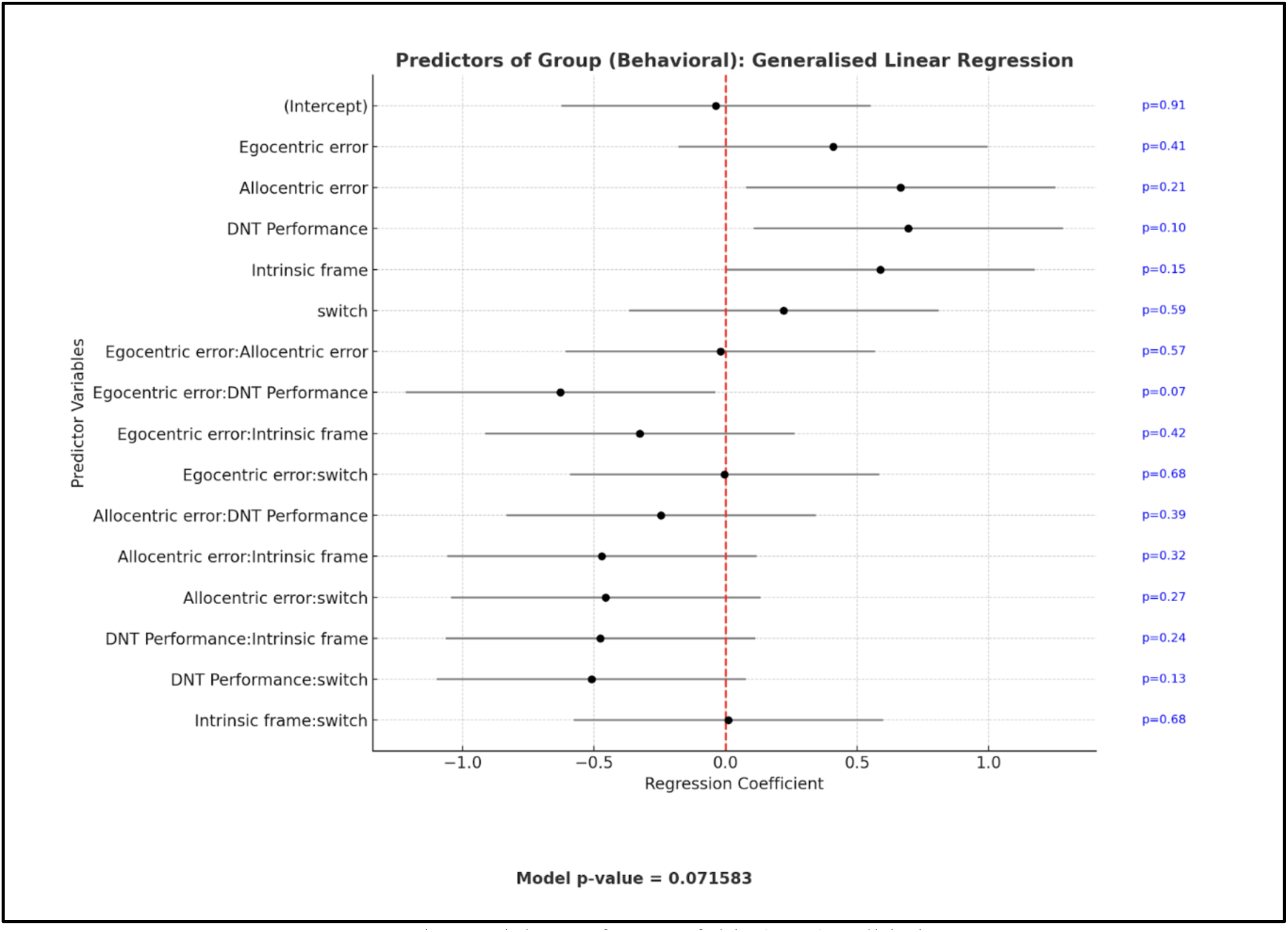
The model stats from K-folds (K=3) validation

